# Unravelling consensus genomic regions conferring leaf rust resistance in wheat via meta-QTL analysis

**DOI:** 10.1101/2021.05.11.443557

**Authors:** Amo Aduragbemi, Jose Miguel Soriano

## Abstract

Leaf rust, caused by the fungus *Puccinia triticina* Erikss (Pt), is a destructive disease affecting wheat and a threat to food security. Developing resistant varieties represents a useful method of disease control, and thus, understanding the genetic basis for leaf rust resistance is required. To this end, a comprehensive bibliographic search for leaf rust resistance quantitative trait loci (QTLs) was performed, and 393 QTLs were collected from 50 QTL mapping studies. Afterwards, a consensus map with a total length of 4567 cM consisting of different types of markers (SSR, DArT, Chip-based SNP markers and SNP markers from GBS) was used for QTL projection, and meta-QTL analysis was performed on 320 QTLs. A total of 75 genetic map positions (gmQTLs) were discovered and refined to 15 high confidence mQTLs (hcmQTLs). The candidate genes discovered within the hcmQTL interval were then checked for differential expression using data from three transcriptome studies, resulting in 92 differentially expressed genes (DEGs). The expression of these genes in various leaf tissues during wheat development was explored. This study provides insight into leaf rust resistance in wheat and thereby provides an avenue for developing resistant varieties by incorporating the most important hcmQTLs.

## Introduction

Leaf rust, caused by the fungal pathogen *Puccinia triticina* Erikss (Pt), causes a significant reduction in grain yield worldwide, and thus, it is considered a disease with significant importance in wheat (*Triticum aestivum* L.) (Roelfs et al. 1992; Khan et al. 2013; Kolmer, 2005). Compared to other fungal rust diseases, such as stem and stripe rust, leaf rust occurs more frequently and has a wider distribution (Bolton et al. 2008). This wide spread of leaf rust may be because the spores of *P. triticina* are transported long distances via wind or humans, thereby causing damage to wheat crops outside their environment or country of origin, as has been observed in various studies in North America (Olmer and Kolmer, 2005; Bolton et al. 2008; Brown and Hovmøll, 2008). The most economical, efficient, environmentally sustainable, and socially acceptable way to manage rust disease globally is to grow rust-resistant varieties (McIntosh et al. 1995; Wiesner-Hanks and Nelson, 2016). Therefore, to understand how best to combine genes and effectively carry out marker-assisted selection (MAS), mapping the target genes conferring resistance in existing parental stocks is essential (Kuchel et al. 2007).

Quantitative trait loci (QTL) mapping is an effective analytical method for studying and manipulating complex traits in crops (Doerge, 2002; Xin et al. 2020). However, QTL mapping has several limitations based on the type of markers, the influence of the environment, the use of different parents, and the size of mapping populations. Therefore, little progress has been made regarding fine mapping and QTL cloning in wheat due to the paucity of high-resolution linkage maps, as the use of simple sequence repeat (SSR) markers is constrained by their inability to saturate the wheat genome (Somers et al. 2004). Consequently, QTLs are frequently present in large genomic regions, and MAS is restricted to small numbers of validated markers (Wang et al. 2015). In recent years, a revolution in the QTL analysis of complex traits has occurred via the introduction of different high-throughput sequencing and genotyping technologies, thus aiding genetic map construction and marker development (Wang et al. 2018a). Numerous QTL studies have been performed in wheat; however, the detected QTLs often do not overlap either in part or in whole due to a different combination of parental lines or studies in multiple environments (Rong et al. 2007). Therefore, there is a need to detect the most promising consensus QTL found among studies using different parents that is stable across environments.

Furthermore, recognizing robust and reliable QTLs and refining their intervals can be achieved by meta-QTL (MQTL) analysis without using expensive resources (Goffinet and Gerber, 2000). The free software BioMercator (Arcade et al. 2004), used for MQTL analysis, allows the compilation of a vast number of genetic maps from various sources and can project QTL to a consensus or reference map (Veyrieras et al. 2007). Thus, MQTL analysis could identify the consensus QTL associated with the trait in multiple environments and genetic backgrounds (Goffinet and Gerber, 2000). QTLs for similar traits may be combined synergistically into MQTL traits by a single MQTL analysis. The reported MQTL method can be used for MAS (Yu et al. 2014; Maccaferri et al. 2015).

Several studies on MQTL-QTL analysis for disease resistance in wheat have been carried out successfully, including those for tan spot resistance (Liu et al. 2020b), Fusarium head blight resistance (Liu et al. 2009; Löffler et al. 2009; Venske et al. 2019; Zheng et al. 2020), leaf rust resistance (Soriano and Royo, 2015), and stem rust resistance (Yu et al. 2014). The first meta-QTL analysis on leaf rust focused only on identifying consensus genomic regions for the trait. In addition, the authors did not use SNP-based genetic maps due to the lack of QTL studies in leaf rust using high-throughput SNP arrays several years ago. However, in this study, we aimed to delve deeper into the genetic architecture underlying leaf rust disease by discovering putative candidate genes from the newly released wheat genome sequence (IWGSC, 2018), incorporating transcriptomic studies, and studying the function of these in genes in different tissues.

## Materials and Methods

### Bibliographic search for wheat leaf rust resistant QTLs

Using Google Scholar (https://scholar.google.com/) and the Web of Science (http://www.webofknowledge.com/), an exhaustive search for publications containing QTLs conferring leaf rust resistance in wheat was performed. For each study, the information collected during the QTL compilation included: (a) the mapping population type: F2:3, recombinant inbred lines (RILs) and double-haploids (DH); (b) nineteen disease resistance traits: disease severity (DS), area under the disease progress curve (AUDPC), leaf infected area (LIA), infection type (IT), infection rate (IR), leaf rust resistance (LR), leaf tip necrosis (LTN), latent period (LP), spore production per unit of sporulating tissue (SPS), spore production per lesion (SPL), infection efficiency (IE), lesion size (LS), early aborted colonies, associated with plant cell necrosis (EA-), early aborted colonies, without plant cell necrosis (EA+), established colonies, associated with plant cell necrosis (EST-), established colonies, without plant cell necrosis (EST+), all necrotic colonies (NEC), maximum disease severity (MDS), and host reaction (HR); (c) number of lines in the mapping population; (d) the LOD (logarithm of the odds) score; (e) the R^2^ value (which denotes the percentage of the phenotypic variation explained (PVE)); and (f) the markers flanking the QTL position (Online Resource 1).

### QTL projection on the consensus map

To project the largest number of QTLs, the high-density consensus map developed by Venske et al. (2019) was used. This consensus map incorporated three marker types: SNPs, DArTs, and SSR markers. The SNPs were sourced from chip-based markers and genotype by sequencing (GBS) (Cavanagh et al. 2013; Saintenac et al. 2013; Wang et al. 2014). SSR markers, including functional markers, were provided from three genetic maps (Wheat, Consensus SSR 2004, Wheat Composite 2004, and Wheat Synthetic x Opata) obtained from the GrainGenes database (https://wheat.pw.usda.gov/GG3/node/976). Wheat consensus map version 4.0, which contains more than 100000 DArTseq markers and nearly 4000 DArT markers developed from over 100 genetic maps, was downloaded from https://www.diversityarrays.com/technology-and-resources/genetic-maps. The initial QTLs were projected following the approach described in Chardon et al. (2004) using BioMercator v4.2 software (Arcade et al. 2004) available at https://urgi.versailles.inra.fr/Tools/BioMercator-V4. Before projecting onto the consensus map, a confidence interval (CI) of 95% was homogenized across the different studies using the following formulas: 530/(N x PVE) for F2:3, 163/(N x PVE) for RILs and 287/(N x PVE) for DH (Darvasi and Soller, 1997; Guo et al. 2006), where N is the number of genotypes in the mapping population, and PVE is the phenotypic variance explained by the QTL.

### Meta-QTL analysis and validation using GWAS

The meta-QTL analysis was conducted using the software BioMercator (Arcade et al. 2004; Sosnowski et al. 2012) by incorporating two approaches for the analysis. The first approach proposed by Goffinet and Gerber (2000) is used when the QTL count for a chromosome is below 10. The second approach proposed by Veyrieras et al. (2007) is used when the QTL count for a chromosome is above 10. For the first approach, the lowest AIC value was selected as the best fit model. However, the best model was selected from the AIC, AICc, AIC3, Bayesian information criterion, and average evidence weight models from the second approach. Therefore, the model with the lowest criteria in at least three of the models was selected and regarded as the best fit model. The MQTL naming convention used by Zheng et al. (2020) was adopted for this study. The MQTLs identified from the meta-QTL analysis were named based on their genetic map positions (gmQTLs). Afterwards, the sequences of the flanking markers of each gmQTL was BLASTed against the Chinese spring reference genome at https://wheat.pw.usda.gov/blast/ (IWGSC, 2018), and their corresponding physical positions were identified, thus resulting in a sequence-based mQTL (smQTL). In addition, the smQTLs found in this study were validated using recent association studies with different genetic backgrounds and environments aimed at identifying loci/QTLs related to leaf rust resistance in wheat.

### Establishment of hcmQTLs and candidate gene mining

To further refine the smQTLs, those with at least five overlapping QTLs having a physical distance lower than 20 Mb and a genetic distance lower than 10 cM were selected and called high confidence mQTLs (hcmQTLs). The annotated reliable genes (HighConfidenceGenesv1.1) within the interval of each hcmQTL were obtained, and their functional annotations were examined at https://wheat-urgi.versailles.inra.fr/Seq-Repository/Annotations.

### Expression of candidate genes within hcmQTL intervals

To check for the candidate genes that were differentially expressed within the hcmQTL intervals, three expression datasets, NCBI-ID ERP013983, SRP041017, and ERP009837 (Dobón et al. 2016; Rudd et al. 2015; Zhang et al. 2014), were used based on experiments reported at ExpVIP (http://www.wheat-expression.com) (Borrill et al. 2016). The ERP013983 dataset consists of differential expression data of ‘Avocet’, a known wheat resistance variety, inoculated with a PST 87/66 strain, with leaf samples collected at 0, 1, 2, 3, and 5 days postinoculation. In the second dataset, SRP041017, the transcriptome of the hexaploid wheat line ‘N9134’ inoculated with the Chinese *Pst* race CYR 31 was compared with the same line inoculated with powdery mildew (*Blumeria graminis* f. sp. *tritici*) (*Bgt*) race E09 at 1, 2, and 3 days postinoculation. The third dataset, ERP009837, consists of differential expression data of the wheat variety ‘Riband’ inoculated with the fungus *Zymoseptoria tritici* (Septoria *tritici* blotch, STB), and the expression data were collected at 1, 4, 9, 14, and 21 days after inoculation. The count data of all the expression data were further analysed, the log2fold change was obtained using the R package Deseq2, and its correction was performed using the R package Apeglm (Zhu et al. 2019). For the DEGs discovered within the refined hcmQTL, Gene Ontology (GO) analysis was performed using the GENEDENOVO cloud platform (https://www.omicshare.com/tools/).

Subsequently, to identify the expression of the reported DEGs in the wheat tissues, the transcriptomics data of ‘Azhurnaya’s 209-sample RNA-sequencing’ project, which examined the developmental timeline of commercial cultivars using a comprehensive array of samples from twenty-four tissue types (Ramírez-González et al. 2018), were used in this study. The stages and their corresponding tissues are as follows: seedling stage: radicle, coleoptile, stem axis, first leaf sheath, first leaf blade, first leaf blade, root, and shoot apical meristem; three leaf stage: third leaf blade and third leaf sheath; tillering stage: first leaf sheath, first leaf blade, shoot axis, and shoot apical meristem; full boot: flag leaf sheath and flag leaf blade; ear emergence: flag leaf sheath and flag leaf blade; anthesis: flag leaf blade night (−0.25 h) 06:45 and fifth leaf blade night (−0.25h) 21:45; milk grain stage: flag leaf sheath, flag leaf blade, and fifth leaf blade (senescence); dough: flag leaf blade (senescence); and ripening: flag leaf blade (senescence). Transcripts per million (TPM) values were used to assess the candidate genes’ level of expression within the hcmQTLs displayed on the heat map using log2 (TPM+1).

## Results

### QTL compilation and projection on the consensus map

A comprehensive search for QTLs conferring resistance to leaf rust resulted in 50 articles published from 1999 to 2020 (Online Resource 1). A total of 393 QTLs widely distributed across the genome were collected (Online Resource 2).

Of the 393 QTLs found, only 320 QTLs had flanking markers and were thus used for MQTL analysis, leaving 73 QTLs with no flanking markers on the consensus map (Online Resource 2). The QTLs were then projected onto the consensus map constructed by Venske et al. (2019). The highest number of QTLs (264) was projected on the B-genome, while the D-genome harboured the lowest number of QTLs (59) (Figure 1). For the A-genome, chromosome 2A had the highest number of projected QTLs, while chromosomes 4A and 7A equally had the lowest (8). Chromosome 1B had the highest number of projected QTLs (41) for the B-genome, while chromosomes 4B and 5B both had the lowest (14). The overall number of QTLs projected on the D-genome was relatively low compared to other genomes, with the highest number of QTLs (22) projected on chromosome 2D, while 6D did not harbour any projected QTLs. When the trait was considered, a large proportion of the projected QTLs were for disease severity (43%), followed by AUDPC (24%) (Figure 1). The rest of the trait categories tagged “others,” comprised 11 traits (SPS, SPL, IE, LS, EA-, EA+, EST-, EST+, NEC, MDS, HR). The PVE varied from 0.01 to 0.97, with 69% of the QTLs reporting a PVE value lower than 0.20. Confidence intervals (CI) ranged from 1.14 cM to 173.11 cM, with an average of 63.98 cM. Most of the QTLs reported a CI lower than 20 cM (81%), with 57% of the QTLs showing a CI lower than 10 cM. Only 2% of the QTLs showed a CI higher than 50 cM.

**Figure 1.**
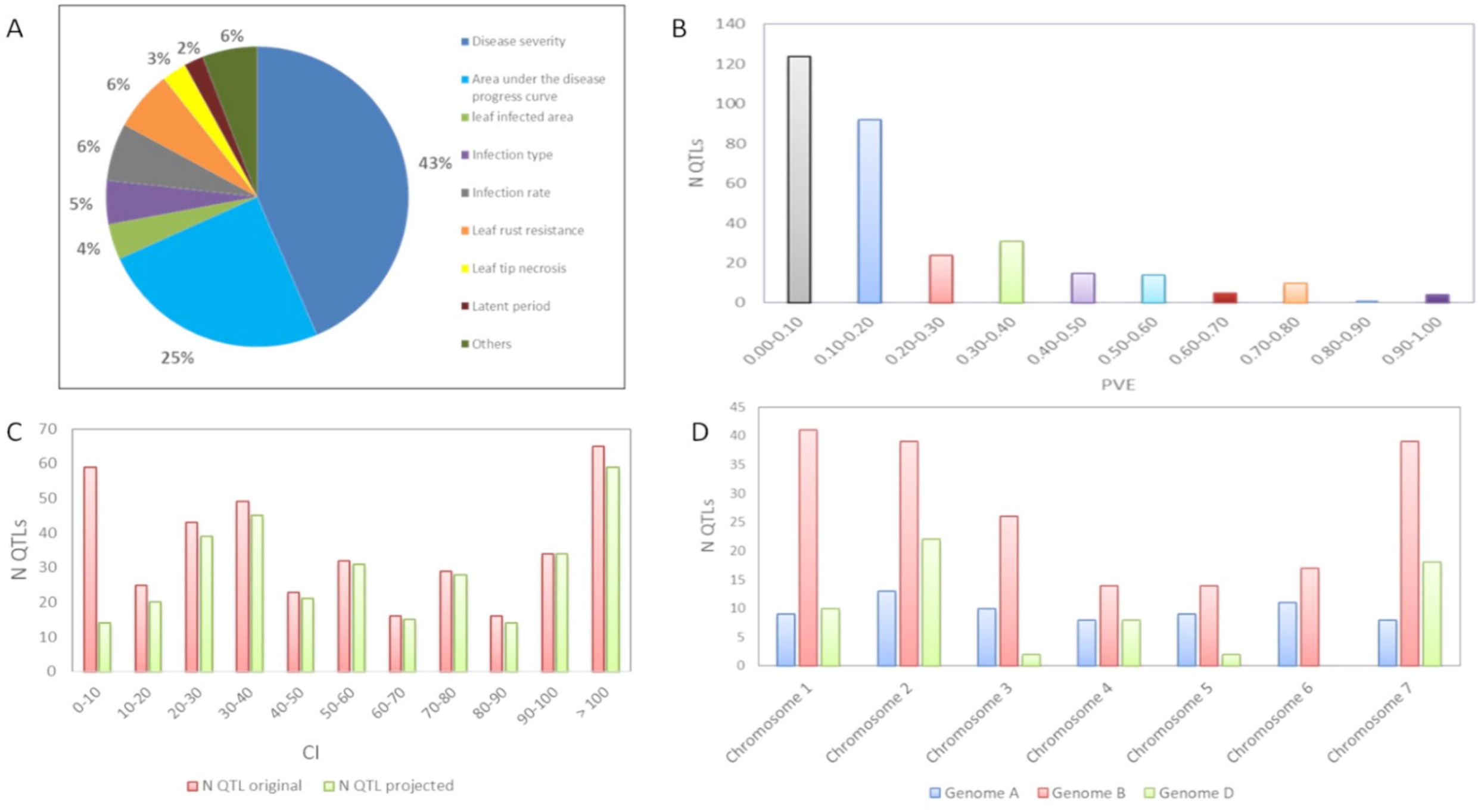
Summary information of QTL projected for MQTL-analysis. Frequency distribution of the number of QTL by (A) Type of resistance trait conferred by the QTL; (B) Phenotypic variation explained (PVE/R^2^); (C) confidence interval; (D) Number of Qtl per chromosome.

### Unravelling consensus regions via MQTL analysis

For the MQTL analysis, the Veyrieras approach was used to analyse all the linkage groups except for chromosomes 1A, 4A, 5A, 7A, 3D, 4D, and 5D because they had fewer than 10 QTLs; thus, the approach of Goffinet and Gerber (2000) was used for their analysis. Overall, 75 gmQTLs were discovered and distributed across all the chromosomes (Figure 1 and Table 1). For the reported gmQTLs, the CI ranged from 0.03 cM to 25.23 cM, with an average of 5.36 cM. For the A genome, 24 gmQTLs were found, with the highest number of gmQTLs (4) on chromosomes 1A, 5A, and 7A, while chromosomes 2A, 3A, 4A, and 6A harboured 3 gmQTLs. For the B genome, a total of 33 gmQTLs were found, representing the genome with the highest number of gmQTLs. Chromosome 2B reported the highest number of gmQTLs (8), followed by 6 gmQTLs on 1B and 6B, and chromosomes 3B and 4B had the lowest number with 3 gmQTLs. For the D genome, a total of 18 gmQTLs were found, with 1D, 2D, and 7D harbouring the most gmQTLs (4), while chromosomes 3D, 4D, and 5D harboured the fewest gmQTLs (2). The physical position of all the gmQTLs was computed, thus yielding a total of 75 smQTLs. The mean physical confidence interval of the smQTLs was 27.47 Mb, which ranged from 0.55 Mb (smQTL1D.1) to 765.3 Mb (smQTL2B.7). The smQTL7B.4 incorporated the highest number of original QTLs. From the smQTLs discovered, the physical interval of six smQTLs was shown to overlap, namely, smQTL1B.2 (636-648 Mb) and smQTL1B.3 (646-662 Mb), smQTL7B.3 (734-744 Mb) and smQTL7B.4 (724-750 Mb), and smQTL5B.4 (634-712 Mb) and smQTL5B.3 (685-704 Mb). Interestingly, none of the smQTLs whose physical intervals overlapped were included in the genetic map.

**Table 1:** The gmQTLs and smQTLs for leaf rust resistance detected in this study

| gm/sm<br>QTL<sup>a</sup> | Chr.<br>b | Pos.<br>(cM)c | CI<br>(cM) d | Flanking markers | CI (Mb) e | N<br>QTL | Traits |
| --- | --- | --- | --- | --- | --- | --- | --- |
| **QTL1A.1** | 1A | 3.0 | 0.0 - 8.8 | 2325584 - BS00026003_51 | 2.9 - 6. | 2 | DS, AUDPC |
| **QTL1A.2** | 1A | 25.2 | 24.2 - 26.3 | wsnp_Ex_c2868_5293115 -<br>1150566 | 7.2 - 10.5 | 2 | AUDPC |
| **QTL1A.3** | 1A | 57.7 | 55.7 - 59.8 | 4909851 - 1151824 | 365.6 - 449.6 | 2 | DS, IR |
| **QTL1A.4** | 1A | 125.6 | 122.6 - 128.6 | wPt-1792 - Xbarc158 | 569.9 - 593.3 | 1 | AUDPC |
| **QTL1B.1** | 1B | 36.4 | 35.4 - 37.3 | 1770278 - 1108935 | 29.6 - 48.1 | 6 | LR, LIA, IT, AUDPC |
| **QTL1B.2** | 1B | 70.3 | 67.5 - 73.2 | Kukri_c18109_331 - 1104505 | 636.3 - 647.6 | 1 | AUDPC |
| **QTL1B.3** | 1B | 79.4 | 78.5 - 80.4 | wsnp_RFL_Contig3951_4390396<br>- 1212781 | 646.2 - 661.6 | 1 | AUDPC |
| **QTL1B.4** | 1B | 91.2 | 90.6 - 91.8 | wsnp_Ra_c53181_56932563 -<br>1103042 | 664.8 - 670.6 | 6 | AUDPC, LR |
| **QTL1B.5** | 1B | 96.9 | 95.1 - 98.8 | BS00000010_51 - 1103838 | 670.8 - 678.6 | 2 | AUDPC, LTN |
| **QTL1B.6** | 1B | 106.8 | 106. - 106.92 | 5971316 -<br>Tdurum_contig49509_606 | 681.0 - 689.0 | 5 | IR, AUDPC, MDS |
| **QTL1D.1** | 1D | 2.6 | 1.5 - 3.8 | wsnp_Ex_c278_538285 -<br>TA003600-0503 | 0.4 - 0.5 | 2 | DS, IR |
| **QTL1D.2** | 1D | 23.9 | 22.7 - 25.0 | 1697264 - Xwmc336 | 1.8 - 10.7 | 5 | DS, IR |
| **QTL1D.3** | 1D | 50.8 | 48.3 - 53.3 | Excalibur_rep_c75168_337 -<br>1004137 | 15.4 - 22.3 | 2 | DS, AUDPC |
| **QTL1D.4** | 1D | 76.1 | 72.8 - 79.3 | 1068907 - BS00028216_51 | 377.9 - 418.3 | 1 | DS |
| **QTL2A.1** | 2A | 15.6 | 13.5 - 17.7 | 1092460 - 3938659 | 71.2 - 104.9 | 1 | IR |
| **QTL2A.2** | 2A | 36.0 | 33.2 - 38.8 | 2263051 - BS00045171_51 | 7.7 - 15.9 | 6 | IR, LP, DS, LS |
| **QTL2A.3** | 2A | 89.9 | 89.7 - 90.1 | 1217255 - 4910857 | 729.9 - 752.3 | 4 | LIA, DS, IR |
| **QTL2B.1** | 2B | 50.9 | 50.2 - 51.6 | IAAV8205 -<br>Excalibur_c1787_1301 | 10.8 - 16.8 | 4 | HR, AUDPC, IR |
| **QTL2B.2** | 2B | 61.0 | 59.7 - 62.4 | BS00022298_51 -<br>wsnp_Ra_c4321_7860456 | 17.6 - 24.9 | 4 | LIA, IT, MDS, AUDPC |
| **QTL2B.3** | 2B | 72.7 | 69.4 - 76.0 | Excalibur_c2454_333 -<br>Excalibur_c13413_1008 | 26.6 - 30.5 | 3 | MDS, AUDPC, 1R |
| **QTL2B.4** | 2B | 96.9 | 94.9 - 98.9 | Tdurum_contig54704_176 - | 58.3 - 65.4 | 5 | AUDPC, LP |
|  |  |  |  | BS00096182_51 |  |  |  |
| **QTL2B.5** | 2B | 106.9 | 106.2 - 107.7 | Ku_c34010_1016 - 1107259 | 154.7 - 176.2 | 4 | IT, AUDPC, LR |
| **QTL2B.6** | 2B | 114.0 | 111.7 - 116.4 | Kukri_c35153_956 - 1187788 | 540.0 - 712.8 | 2 | LR |
| **QTL2B.7** | 2B | 128.1 | 126.2 - 130.1 | 1094755 - 1381445 | 737.2 - 765.3 | 2 | AUDPC, LIA |
| **QTL2D.1** | 2D | 2.7 | 1.0 - 4.4 | IAAV298 - D_contig74612_253 | 1.7 - 10.3 | 7 | DS, LP |
| **QTL2D.2** | 2D | 33.4 | 31.6 - 35.3 | 1090614 - 1059345 | 37.2 - 49.9 | 5 | DS, SPS, LTN |
| **QTL2D.3** | 2D | 50.6 | 48.8 - 52.4 | D_GBB4FNX01DJHXL_88 - 1101216 | 50.9 - 75.7 | 1 | DS |
| **QTL2D.4** | 2D | 72.0 | 69.7 - 74.3 | RAC875_c42714_775 - 1102692 | 461.3 - 565.0 | 4 | DS |
| **QTL3A.1** | 3A | 11.7 | 8.3 - 15.1 | 1334535 - 4394604 | 1.3 - 13.3 | 1 | DS |
| **QTL3A.2** | 3A | 23.9 | 21.5 - 26.4 | 1243077 - wsnp_Ex_c8409_14170476 | 21.2 - 24.6 | 2 | DS, AUDPC |
| **QTL3A.3** | 3A | 70.8 | 70.6 - 71.0 | RAC875_c583_341 - IACX333 | 662.4 - 674.7 | 8 | LIA, DS, IT, AUDPC |
| **QTL3B.1** | 3B | 11.7 | 10.7 - 12.7 | wsnp_Ex_c30368_39293103 - 1213030 | 3.4 - 6.4 | 3 | LIA, IT |
| **QTL3B.2** | 3B | 27.2 | 24.6 - 29.7 | 1230466 - 1121919 | 11.7 - 24.4 | 8 | AUDPC, SPL, IT, LP, IR, IE, LS |
| **QTL3B.3** | 3B | 70.1 | 69.3 - 71.0 | 3947488 - 1262486 | 580.5 - 615.8 | 5 | AUDPC, IT, LIA |
| **QTL3D.1** | 3D | 9.5 | 6.7 - 12.2 | 1228026 - 2244235 | 7.3 - 13.3 | 1 | DS |
| **QTL3D.2** | 3D | 21.3 | 11.8 - 30.9 | 3026734 - Kukri_c5252_107 | 12.9 - 31.7 | 1 | DS |
| **QTL4A.1** | 4A | 27.1 | 21.6 - 32.7 | 4909758 - 3022955 | 56.3 - 91.7 | 2 | IT, DS |
| **QTL4A.2** | 4A | 38.6 | 35.5 - 41.6 | 3948059 - 3021557 | 108.8 - 177.5 | 3 | LR, DS, IR |
| **QTL4A.3** | 4A | 54.6 | 50.9 - 58.2 | BobWhite_c12128_187 - 1387412 | 610.5 - 670.9 | 3 | LR, DS |
| **QTL4B.1** | 4B | 34.1 | 32.4 - 35.7 | 1105735 - wsnp_Ex_c30695_39579408 | 14.3 - 20.6 | 5 | DS, IR |
| **QTL4B.2** | 4B | 50.5 | 48.6 - 52.4 | 1210331 - Tdurum_contig4974_1386 | 53.0 - 95.7 | 5 | DS |
| **QTL4B.3** | 4B | 99.5 | 96.1 - 102.9 | 1211581 - BS00016834_51 | 656.9 - 667.7 | 2 | IR, DS |
| **QTL4D.1** | 4D | 36.7 | 35.4 - 37.9 | 2261222 - 1201923 | 15.0 - 23.3 | 4 | IT, LIA, AUDPC, DS |
| **QTL4D.2** | 4D | 78.4 | 72.8 - 84.1 | 1314743 - 1130355 | 465.2 - 497.9 | 4 | DS, AUDPC, LS, SPL |
| **QTL5A.1** | 5A | 30.2 | 27.4 - 33.1 | 3064538 - 995502 | 26.8 - 34.7 | 1 | LTN |
| **QTL5A.2** | 5A | 41.3 | 39.1 - 43.5 | BS00022110_51 - | 440.3 - 549.6 | 3 | DS, IR |
|  |  |  |  | wsnp_Ra_c17216_26044790 |  |  |  |
| **QTL5A.3** | 5A | 55.0 | 52.2 - 57.8 | 3025345 - 1215538 | 554.3 - 586.3 | 4 | DS, AUDPC |
| **QTL5A.4** | 5A | 130.7 | 124.1 - 137.4 | Xwmc110 - 1084271 | 644.9 - 685.0 | 1 | EA- |
| **QTL5B.1** | 5B | 18.9 | 15.2 - 22.6 | wsnp_Ex_rep_c105478_8989163<br>4 - 3021398 | 485.9 - 531.2 | 3 | AUDPC, LR |
| **QTL5B.2** | 5B | 46.1 | 44.1 - 48.2 | 3023016 - 982446 | 609.1 - 657.4 | 1 | IT |
| **QTL5B.3** | 5B | 69.2 | 66.4 - 72.0 | 2293625 - 1060195 | 685.2 - 704.2 | 4 | LIA, IT, LTN, AUDPC |
| **QTL5B.4** | 5B | 142.1 | 135.9 - 148.4 | Xwmc235 - 1280229 | 634.2 - 711.9 | 4 | IT, LIA |
| **QTL5D.1** | 5D | 8.15 | 0.0 - 25.2 | 1091611 - 2289853 | 1.6 - 3.5 | 1 | DS |
| **QTL5D.2** | 5D | 43.4 | 41.8 - 44.9 | 1038800 -<br>BobWhite_c29857_228 | 5.6 - 25.3 | 1 | DS |
| **QTL6A.1** | 6A | 15.5 | 13.3 - 17.8 | Kukri_c38982_165 - 1112318 | 11.1 - 17.9 | 4 | DS, LP |
| **QTL6A.2** | 6A | 33.5 | 30.9 - 36.1 | 4909358 - 1238725 | 33.5 - 49.8 | 5 | DS, AUDPC, LR |
| **QTL6A.3** | 6A | 48.2 | 44.2 - 52.1 | 1237185 -<br>wsnp_Ku_c4296_7807837 | 447.4 - 520.7 | 2 | DS, LR |
| **QTL6B.1** | 6B | 6.7 | 6.3 - 7.1 | 1139996 - GENE-4226_445 | 4.6 - 9.0 | 2 | DS, AUDPC |
| **QTL6B.2** | 6B | 19.6 | 14.2 - 25.1 | RAC875_c62256_577 -<br>wsnp_Ex_c4815_8597064 | 17.7 - 30.1 | 2 | DS, AUDPC |
| **QTL6B.3** | 6B | 44. | 40.5 - 48.5 | IAAV8279 -<br>wsnp_Ex_c19874_28891457 | 514.0 - 665.1 | 4 | DS, AUDPC |
| **QTL6B.4** | 6B | 71.3 | 70.3 - 72.2 | 1094048 - 1228581 | 701.4 - 708.1 | 2 | DS, AUDPC |
| **QTL6B.5** | 6B | 82.8 | 81.2 - 84.4 | 3949402 - 2275538 | 713.6 - 715.2 | 4 | DS, AUDPC, LR |
| **QTL6B.6** | 6B | 106.2 | 105.9 - 106.4 | 1231000 -<br>Tdurum_contig10729_64 | 717.9 - 721.0 | 2 | DS, AUDPC |
| **QTL7A.1** | 7A | 74.4 | 73.8 - 74.9 | 1099858 - BS00047811_51 | 69.7 - 79.6 | 1 | LR |
| **QTL7A.2** | 7A | 86.9 | 80.0 - 93.9 | 7345647 - IACX2471 | 554.4 - 669.7 | 1 | LR |
| **QTL7A.3** | 7A | 117.0 | 107.7 - 126.3 | Tdurum_contig29834_165 -<br>1002615 | 679.9 - 714.6 | 1 | DS |
| **QTL7A.4** | 7A | 130.2 | 128.9 - 131.4 | 3956725 - wPt-0745 | 716.4 - 736.2 | 5 | DS, AUDPC |
| **QTL7B.1** | 7B | 74.9 | 73.0 - 76.8 | Tdurum_contig47317_100 -<br>wsnp_Ex_c19_39269 | 675.1 - 693.3 | 7 | IR, LR, AUDPC, IT, LTN |
| **QTL7B.2** | 7B | 93.5 | 92.3 - 94.7 | 1114524 - BobWhite_c2892_167 | 700.7 - 721.2 | 3 | LR, LP, AUDPC |
| **QTL7B.3** | 7B | 122.3 | 120.4 - 124.3 | wPt-2933 - Xwmc10 | 734.3 - 744.9 | 3 | LR, LTN, IR |
| **QTL7B.4** | 7B | 141.7 | 141.7 - 141.7 | Xbarc32 - RAC875_c525_1372 | 723.9 - 750.1 | 9 | EA-, EA+, EST-, LIA,<br>EST+, NEC, LP, AUDPC |
| **QTL7D.1** | 7D | 25.8 | 25.4 - 26.2 | D_contig24676_377 - 1103360 | 167.6 - 185.1 | 8 | LTN, DS, LR, AUDPC, LS |
| **QTL7D.2** | 7D | 42.4 | 41.2 - 43.6 | 2246156 - 2244056 | 569.2 - 591.7 | 4 | MDS, AUDPC, DS |
| **QTL7D.3** | 7D | 58.5 | 57.9 - 59.1 | 1242088 - 1123852 | 610.9 - 629.7 | 4 | AUDPC, DS, LTN |
| **QTL7D.4** | 7D | 148.6 | 137.7 - 159.5 | 3533423 - wPt-666144 | 636.3 - 638.2 | 1 | LTN |

<sup>a</sup> gmQTL, genetic map positions; smQTL, sequence-based mQTL (smQTL)
<sup>b</sup> Chromosomes
<sup>c</sup> Genetic position
<sup>d</sup> Genetic position confidence interval
<sup>e</sup> physical position confidence interval

### Association study validation and colocalization with leaf rust genes

The loci resistant to leaf rust across different genetic backgrounds and environments identified in recent association studies (Online Resource 3) were used to validate the MQTLs discovered in this study. A total of 51 MTAs identified were colocalized with 29 MQTLs; thus, some MQTLs integrated more than one MTA (Online Resource 4, Figure 2). Furthermore, eight leaf rust genes colocalized with some smQTLs found in this study (Online Resource 5, Figure 2), such as smQTL1D.2 colocalizing with Lr60 and Lr42 and smQTL7B.3 colocalizing with the Lr68 and Lr14a genes. Additionally, smQTL1B.4 and smQTL2B.5 both colocalized with two leaf rust genes, Lr46 and Lr13, respectively, conferring adult plant resistance.

**Figure 2.**
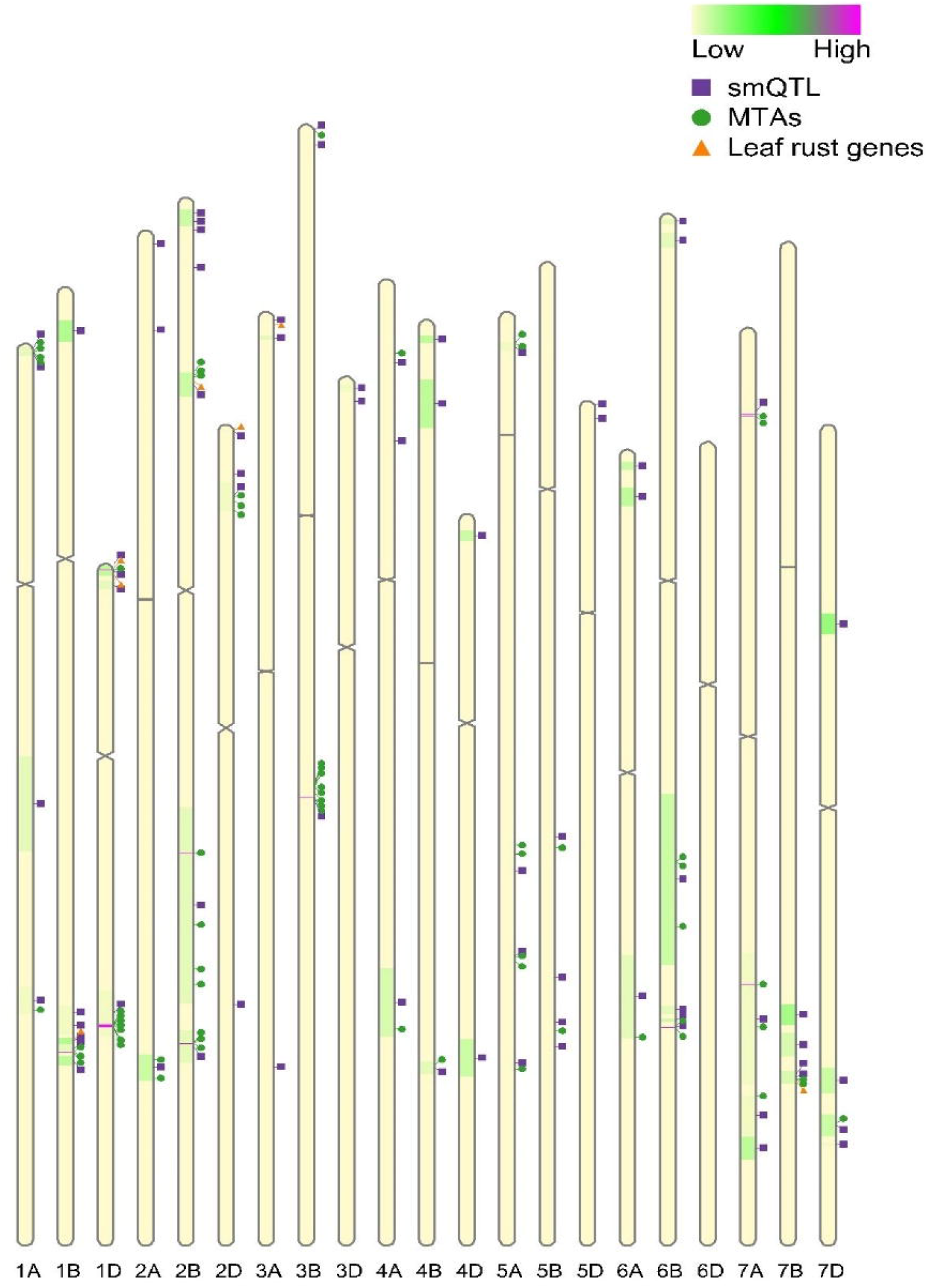
Distribution of the smQTLs, MTAs, and co-localized leaf rust genes with reference to the Chinese spring genome. The chromosome coloration correlates to the number of initial QTLs: the lighter color connotes less QTL. The centromere of each chromosome is represented by the constriction.

### Candidate gene mining of established high confidence mQTLs (hcmQTLs)

To further improve the quality of the smQTLs discovered, they were further refined to regions termed high confidence mQTLs (hcmQTLs). The hcmQTLs consist of 15 consensus regions (Online Resource 6), having an average confidence interval and physical interval of 2.9 cM and 12.04 Mb, respectively. Overall, each hcmQTL cluster contained at least five QTLs. The B genome had the highest number of these hcmQTLs (7). Within the B genome, chromosome 1B contained the highest number of hcmQTLs (3). In addition, hcmQTL4B.1 had the smallest physical interval (6.25 Mb), while hcmQTL7A.4 had the largest interval, covering 19.78 Mb. Afterwards, candidate gene mining within hcmQTLs revealed 2240 genes, with hcmQTL7A.4 possessing the highest number of candidate genes and hcmQTL2B.4 possessing the lowest number (18) of candidate genes.

### Discovering differentially expressed genes (DEGs) within hcmQTLs

Owing to the lack of transcriptomic data repositories for leaf rust in wheat, we decided to use transcriptomic expression data for three fungal diseases in wheat, stripe rust, powdery mildew, and *Septoria tritici* blotch, to explore the DEGs within these hcmQTLs. The ERP013983 dataset revealed 541 DEGs with 221 downregulated genes, 255 upregulated genes, and 65 genes that were downregulated under several conditions and upregulated under others (Online Resource 7, Figure 3). From this dataset, hcmQTL7A.4 had the highest number of DEGs (59), while hcmQTL2B.4 had the lowest (12). The SRP041017 dataset revealed 289 DEGs, where 131 genes were downregulated, 154 genes were upregulated, and 4 genes were downregulated at one time point and upregulated in others (Online Resource 8, Figure 3). From this dataset, hcmQTL2A.2 had the highest number of DEGs (39), while hcmQTL2B.4 and hcmQTL4B.1 (4) had the fewest DEGs (4). From the third dataset, ERP009837, a total of 327 DEGs were discovered, with 125 genes downregulated, 198 genes upregulated, and 4 genes downregulated at one time point and upregulated in others (Online Resource 9, Figure 3). From this dataset, hcmQTL2A.2 had the highest number of DEGs (34), while hcmQTL2B.4 had the lowest number of DEGs (3). A total of 92 genes were found to be differentially expressed across the three expression datasets used. The 92 DEGs within the hcmQTL interval were analysed for Gene Ontology (GO) enrichment (Table 2). The most significantly enriched GO terms associated with biological processes were for metabolic (44 genes) and cellular processes (31 genes) (Online Resource 10, Figure 4). The most significantly enriched GO terms associated with molecular function were for catalytic activities (41 genes) and binding (39 genes). In terms of cellular components, the genes were enriched mainly in the cell membrane and its components.

**Figure 3.**
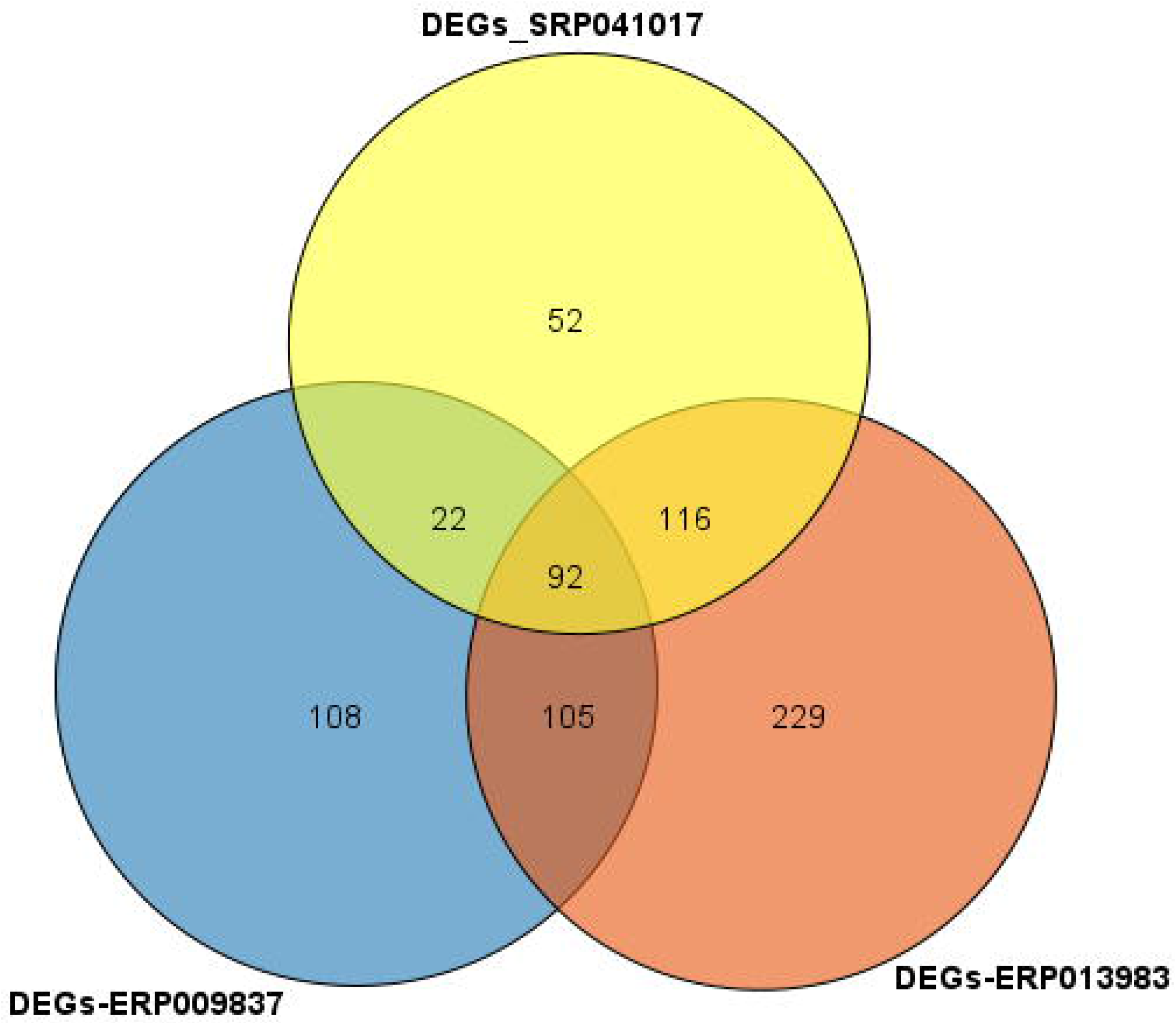
Venn diagram depicting the number of differentially expressed genes (DEGs) from three transcriptomic data sets. The Venn diagram visually illustrates the number of differentially expressed genes (DEGs) that are identified in the various transcriptomic data sets (A) dataset for stripe rust (B) dataset for three fungi diseases in wheat: fusarium head blight, powdery mildew, and Septoria tritici blotch.

**Table 2:**
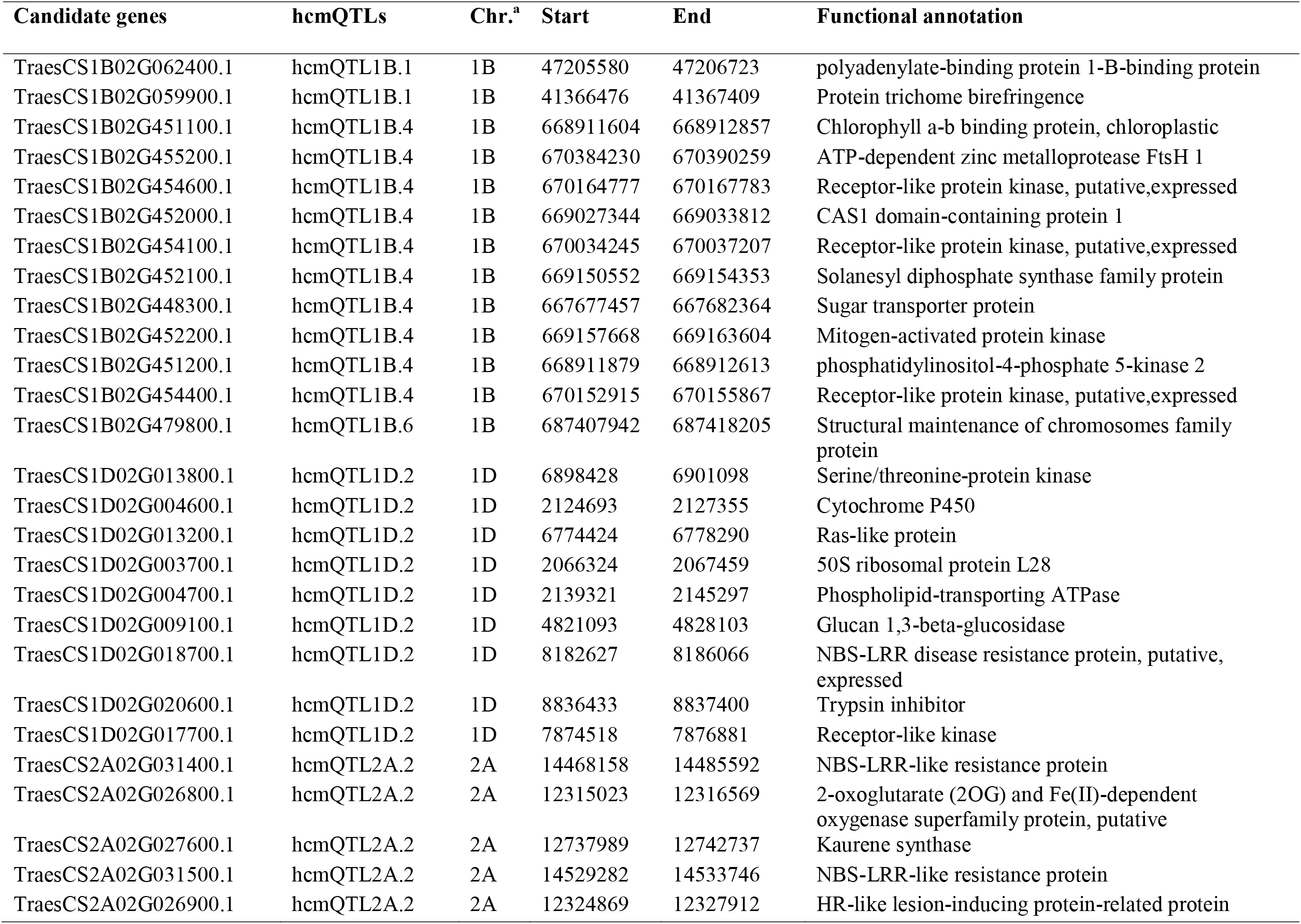
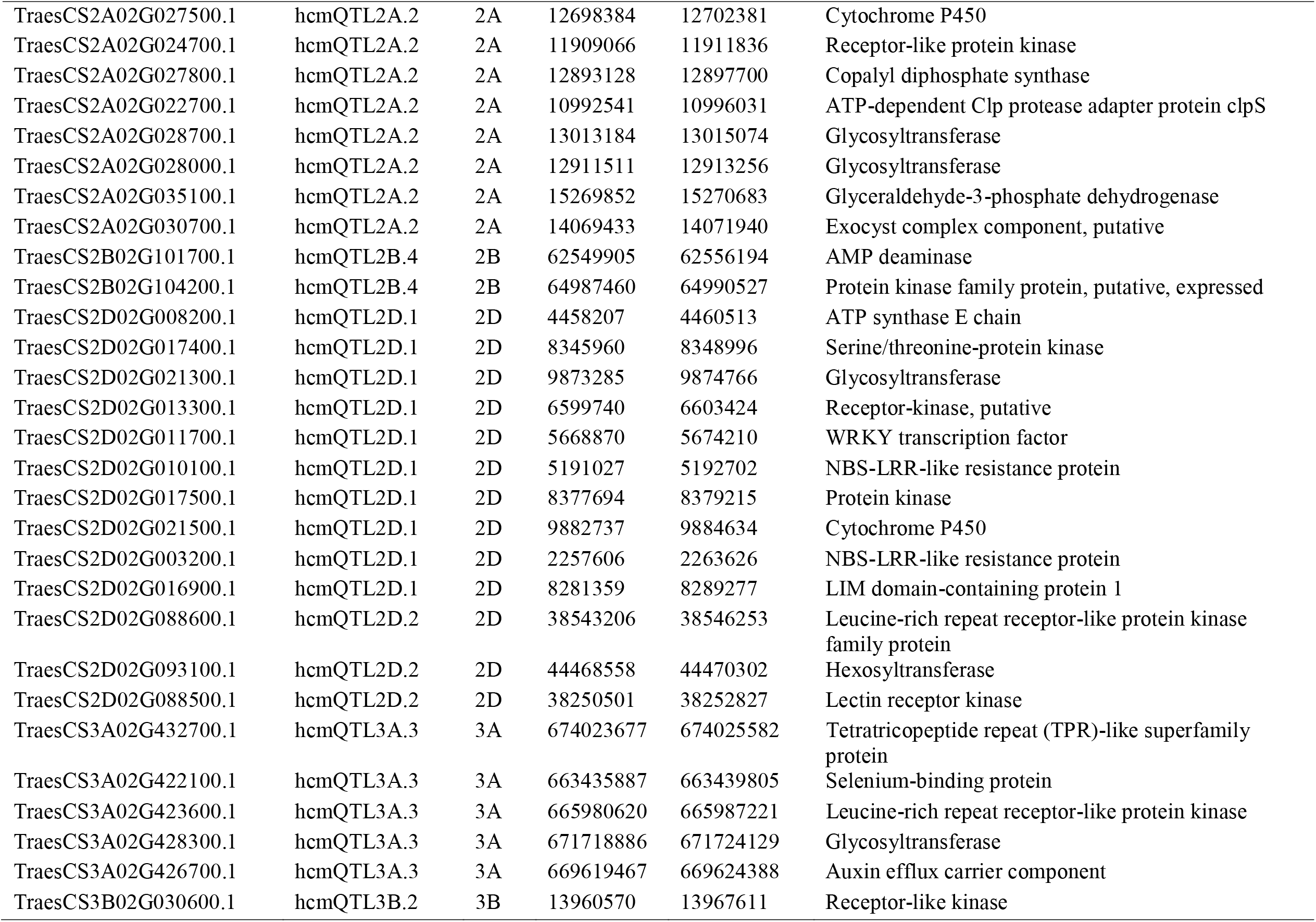
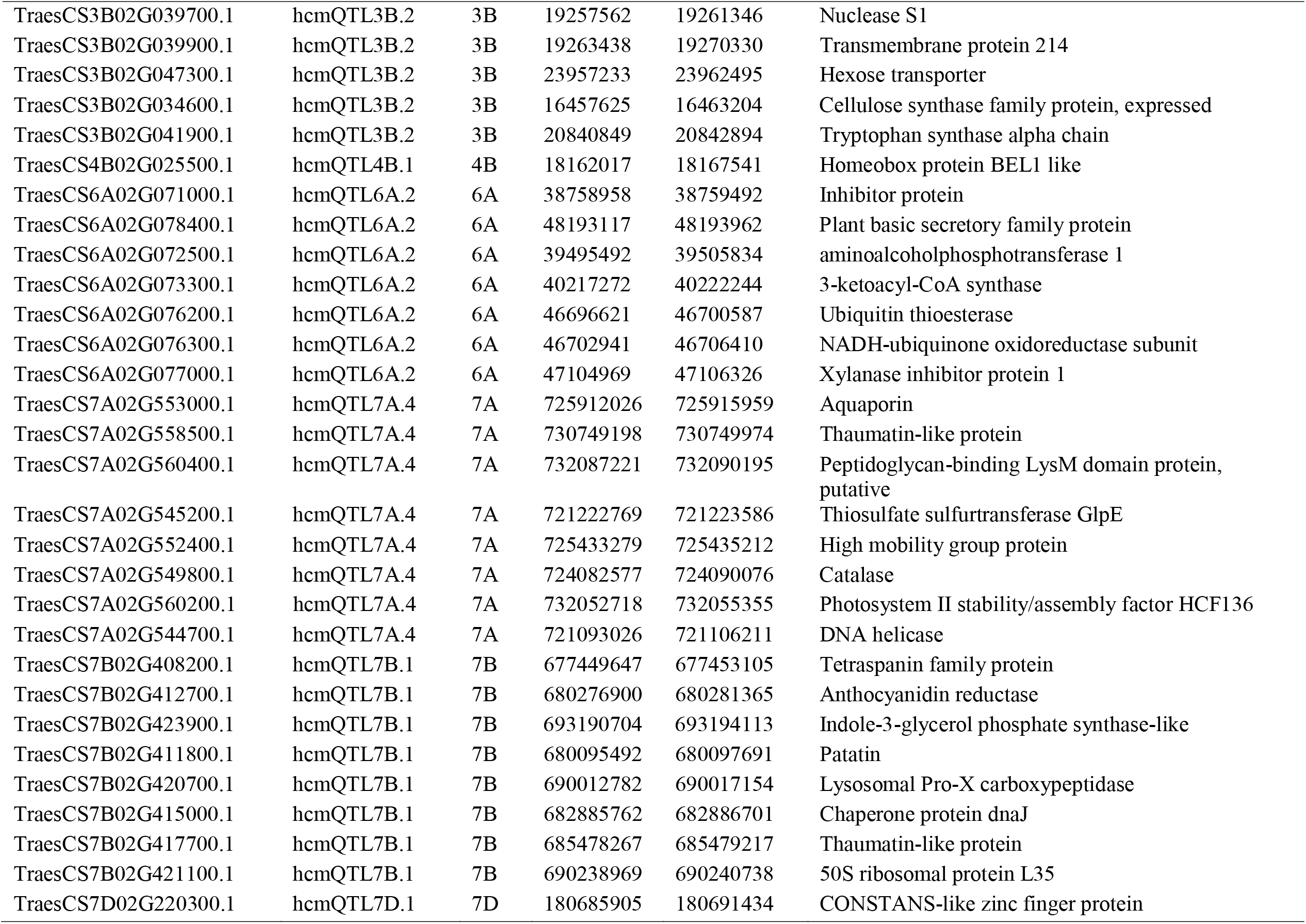
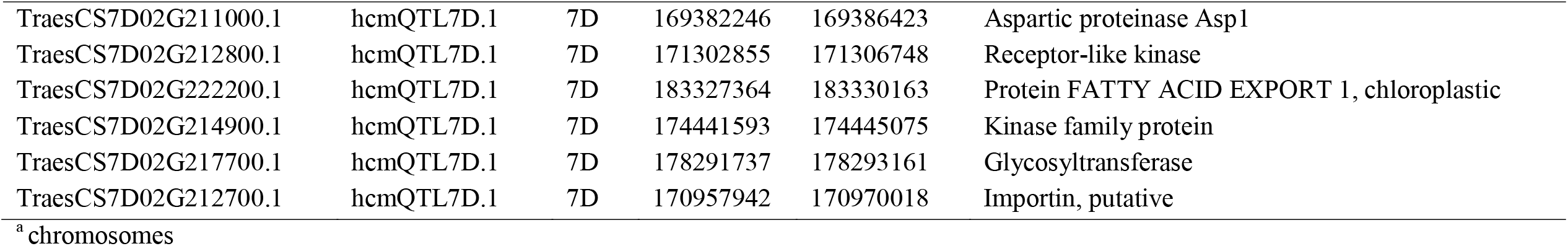
High confidence MQTLs that showed differential expression across the three transcriptomics datasets.

**Figure 4.**
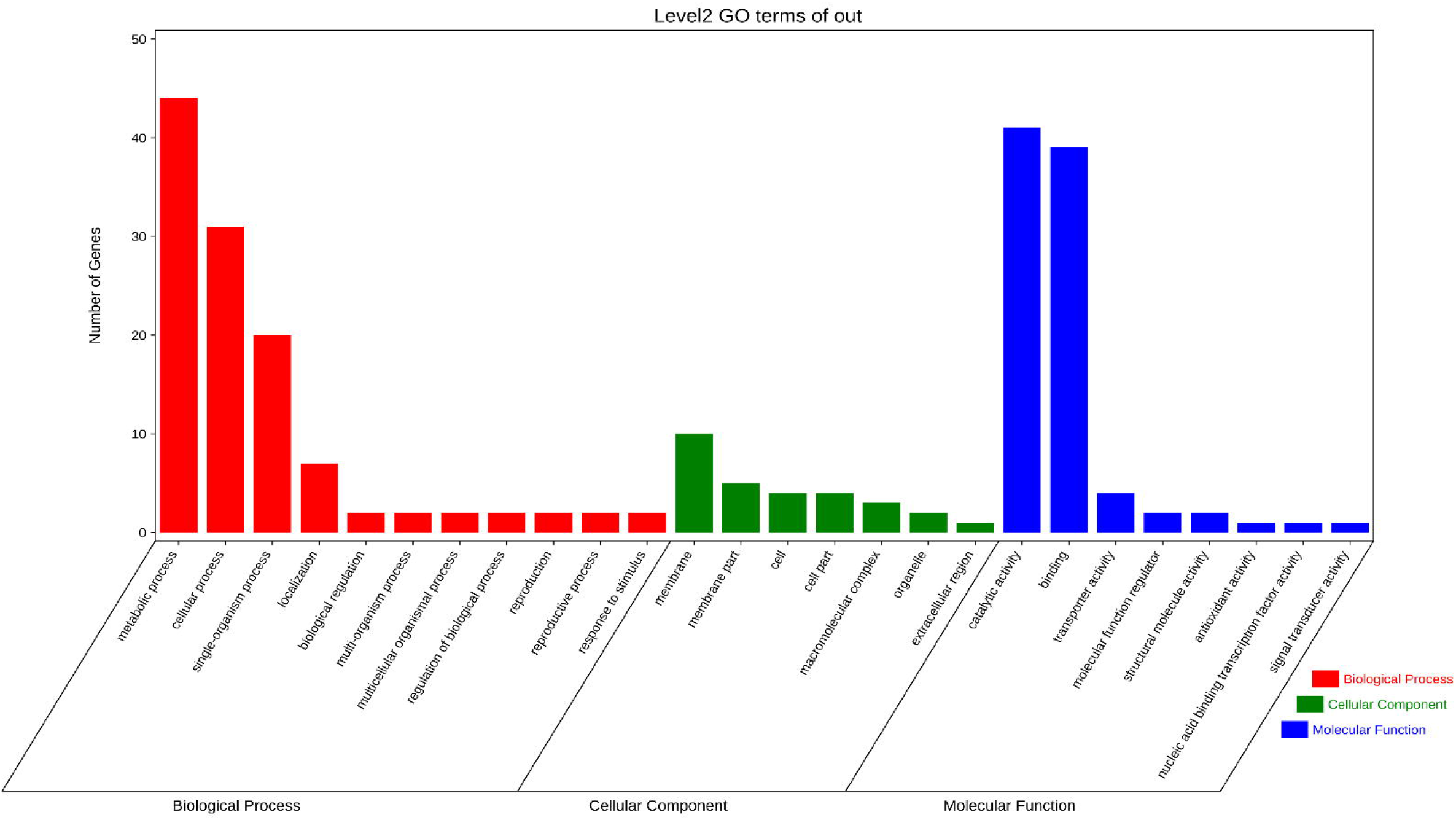
Level 2 GO terms for DEGs in the hcmQTL regions.

### Tissue-specific expression profile of DEGs within hcmQTLs

To understand the expression of the DEGs within the hcmQTL intervals across different tissues at different wheat development stages, we used DEGs expressed in all the transcriptomics datasets SRP041017, ERP013983, and ERP009837. Row clustering was applied, and as a result, the 92 DEGs fell into two classes (I-II) based on their expression patterns (Figure 5). Genes in class I showed moderate to high expression in the flag leaf blade and fifth leaf blade at the anthesis stage when compared to other stages of growth. Moreover, for class II, more genes were highly expressed in the first leaf sheath at the tillering stage. The DEGs in class II accounted for more than half of the overall DEGs, and they showed contrasting expression patterns to those shown by genes in class I. For class I, at the seedling stage, the genes TraesCS1D02G003700 (hcmQTL1D.2) and TraesCS7B02G421100 (hcmQTL7B.1) showed moderate expression in the stem axis, while at the adult stage, TraesCS6A02G072500 (hcmQTL6A.2) was highly expressed in the fifth leaf blade at the anthesis stage, and TraesCS1D02G017700 (hcmQTL1D.2) showed high expression in the flag leaf at the dough stage. Furthermore, at the seedling stage, more class II genes were moderately expressed in the radicle and roots, with only TraesCS7D02G211000 (hcmQTL7D.1) showing expression in the shoot apical meristem. At the adult stage, the gene TraesCS7D02G220300 (hcmQTL7D.1) was highly expressed in the fifth leaf blade at the anthesis stage.

**Figure 5.**
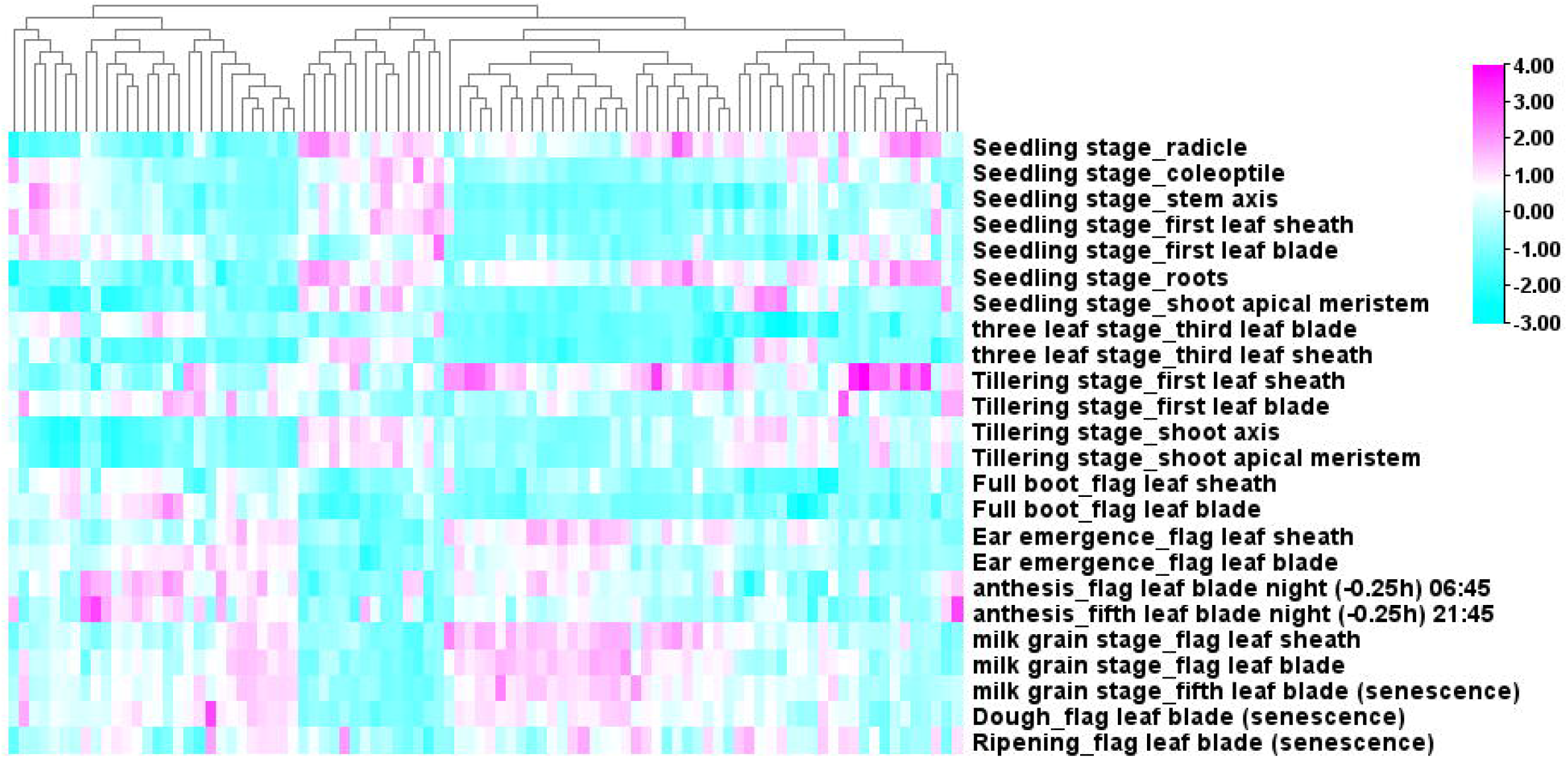
Expression pattern of 90 candidate genes in 24 tissues. All transcriptome data was downloaded from expVIP (http://www.wheat-expression.com). The changes in color from sky blue to pink signifies alteration in level of expression from low to high.

## Discussion

### Establishment of gmQTL and smQTL regions

To gain deeper insight into the control of leaf rust resistance in wheat, a meta-QTL analysis was performed based on the numerous QTLs conferring leaf rust resistance identified in the literature from various independent studies. The first step to identifying consensus regions via meta-QTL analysis is the projection of the original QTLs onto a consensus or reference map.

A feature of the consensus map and QTL database was that the B genome reported the highest marker saturation, and thus, the highest number of QTLs was mapped to this genome, which is in agreement with previous studies characterizing genetic diversity and unravelling complex traits for disease resistance in bread wheat (Li et al. 2015; Soriano and Royo, 2015; Wang et al. 2014). The D genome presented a lower number of QTLs, as previously found in other meta-QTL analyses for disease resistance in wheat (Soriano and Royo, 2015; Venske et al. 2019; Liu et al. 2020b; Zheng et al. 2020). Furthermore, no QTLs were found on chromosome 6D, as discovered in previous meta-QTL analysis studies on leaf rust and Fusarium head blight diseases in wheat (Soriano and Royo, 2015; Zheng et al. 2020). A possible explanation for the limited QTLs located on the D genome across various disease studies could be the low level of polymorphism associated with the D genome. In this study, a larger number (81.4%) of QTLs was projected onto the consensus map compared to the fewer number of QTLs projected in a previous meta-QTL analysis for leaf rust (Soriano ad Royo, 2015) (44%). A possible reason could be due to the different consensus maps used. In this study, we used a high-density consensus map that combined SSRs and markers obtained from high-throughput genotyping platforms, in contrast to the consensus map used in the previous study from Soriano and Royo (2015). Consequently, the number of genomic regions (gmQTLs) discovered in this study was higher than those reported in Soriano and Royo (2015), at 75 and 48, respectively. For the gmQTLs discovered in this study, the confidence interval ranged from 0.03 cM to 25.23 cM, which was significantly reduced by more than 50% from the confidence interval of the original QTLs, ranging from 1.14 cM to 173.11 cM. In addition, in the present study, the physical position of the gmQTLs was reported due to the release of the wheat genome sequence (IWGSC, 2018), improving the mapping resolution of the genome regions (smQTLs) and helping the identification of candidate genes. These analyses enhance the results provided by studies published prior to the release of the genome sequence. In this study, we discovered 7 smQTLs incorporating at least five original QTLs and having a confidence interval of less than 10 Mb, making them the most promising for candidate gene identification.

### Colocalization of smQTLs with leaf rust resistance genes and traits

To strengthen the location of smQTLs discovered in this study, a search for colocalization of leaf rust resistance genes and smQTLs was performed. More than 61 leaf rust genes have been mapped and documented in wheat (Kim et al. 2020), and 4 of them have been cloned (Hafeez et al. 2021). A total of six leaf rust genes (Lr13, Lr14a, Lr46, Lr68, Lr63, Lr60, Lr42, and Lr41) were found to colocalize with smQTLs. Interestingly, the colocalization of Lr13, Lr14a, and Lr46 with smQTL2B.5, smQTL7B.3, and smQTL1B.4 on chromosomes 2B, 7B, and 1B, respectively, in this study was in agreement with the results obtained by Soriano and Royo (2015). As reported by these authors, Lr68 was found to have a tight association with MQTL33 (colocalized with Lr14a) on chromosome 7B; however, in this study, smQTL7B.3 colocalized with both leaf rust genes (Lr14a and Lr68), thus confirming the usefulness of using highly saturated consensus maps for meta-QTL analysis. The gene Lr14a, known to confer adult plant resistance, is thought to have evolved from emmer wheat cv. Yaroslav (McFadden, 1930) and is associated with the stem rust and powdery mildew resistance genes Sr17 and Pm5. Lr68, on the other hand, confers seedling resistance to the majority of *P. triticina* isolates with low to medium infection types and is linked to small but noticeable leaf tip necrosis (Herrera-Foessel et al. 2012).

Consequently, the smQTL7B.3 region not only confers seedling and adult plant resistance to leaf rust but also constitutes a region of multiple disease resistance. Additionally, smQTL1D.2 colocalized with two leaf rust resistance genes (Lr60 and Lr42). Hiebert et al. (2008) found that Lr60 is 13.5 cm distal to Lr21, which would position Lr60 and Lr42 approximately 40 cm apart (Huang et al. 2003; Somers et al. 2004). The association between Lr60 and Lr42 has not been confirmed, but in this study, we discovered that both genes were located in the same smQTL, with a confidence interval of 8.8 Mb. This supports possible linkage between the two genes; however, a genetic linkage test needs to be carried out to corroborate this claim. Furthermore, Lr60 is known to confer seedling resistance, while Lr42 confers adult plant resistance. Thus, the smQTL1D.2 region has the potential to provide more qualitative resistance against leaf rust. Additionally, smQTL1B.4 colocalized with Lr46, a gene known to increase the latent period and reduce the frequency of infection and uredinial size in a similar manner to Lr34 (Drijepondt and Pretorius, 1989; William et al. 2003). There is also a tight linkage between Lr46 and a stripe rust gene (*Yr29)*, which is similar to the linkage between Lr34 and Yr18 (McIntosh, 1992; Singh 1992). Consequently, the smQTL1D.2 region confers resistance to both leaf and stripe rust in wheat, thus making this region a hotspot for selecting multiple disease resistance in wheat.

Most of the smQTLs discovered in this study clustered QTLs conferring two or more resistance traits. This phenomenon was also discovered by Kolmer et al. (2018) using RILs, suggesting that they experienced higher disease severity levels and leaf rust responses. In another study, Ren et al. (2012) also discovered that maximum disease severity had a significant association with the area under the disease progress curve (AUDPC) across diverse environments, and this finding was in agreement with previous studies (Wang et al. 2005b; Liang et al. 2006; Lan et al. 2009). Consequently, this result indicates the possibility of replacing AUDPC with MDS. A possible explanation for this could be that when two or more traits are mapped to the same region, they are most likely under the same genetic control, as suggested by Lu et al. (2017). Furthermore, effects arising from tight linkage and pleiotropism could also be a possible explanation.

### Candidate genes within hcmQTLs and their role in leaf rust resistance

The search for candidate genes was extended to hcmQTLs within 20 Mb, thus yielding 15 hcmQTLs. hcmQTLs also have a small confidence interval compared to smQTLs, thus making them more reliable and useful for QTL selection in breeding programmes. The hcmQTLs harboured a total of 2240 genes, after which 92 DEGs were narrowed down. Two main types of disease resistance are used in breeding programmes: seedling resistance and adult plant resistance. Thus, the analysis of the expression of the candidate genes across different tissues and developmental stages can inform us of their potential role in seedling or adult plant resistance. Five out of the 92 genes expressed across the three transcriptomic data sets, TraesCS7D02G212800, TraesCS6A02G073300, TraesCS2B02G104200, TraesCS1D02G003700, and TraesCS2D02G021300, showed moderate expression in the first leaf sheath at the seedling stage. TraesCS7D02G212800 and TraesCS2B02G104200 encode a receptor-like kinase (RLK) and protein kinase family protein, respectively, and both proteins play a crucial role in contributing to disease resistance in wheat. Plant protein kinases, as well as receptor-like kinases, govern the detection and activation of diverse developmental and physiological signals, particularly those involved in defence and symbiosis (Rentel et al. 2004; AbuQamar et al. 2008; Fu et al. 2009; Garcia et al. 2012). Prior studies found that various RLK genes coding wheat leaf rust kinases (WLRKs) were conserved in wheat, with the most studied member of the WLRK family being LRK10, which is genetically linked to the Lr10 locus (Feuillet *et al*. 1997, 1998, 2001). Gu et al. (2020), in a recent study, uncovered an RLK gene that plays an important role in resistance to *P. triticina* infection and has a positive regulatory effect on the hypersensitive reaction (HR) cell death process induced by *P. triticina*. TraesCS6A02G073300, encoding a 3-ketoacyl-CoA synthase, has been reported to harbour quantitative trait nucleotides (QTNs) in close proximity to leaf rust genes in wheat (Fatima et al. 2020). The 50S ribosomal protein L28 encoded by TraesCS1D02G003700 belongs to the ribosomal protein family, and members of this family have been shown to confer tolerance against fungal pathogens in plants (Yang et al. 2013). Furthermore, TraesCS7D02G217700, encoding a glycosyltransferase, was highly and moderately expressed in the first leaf blade and leaf sheath, respectively, at the seedling stage. According to Bolton et al. (2008), two pathogen-responsive genes encoding glycosyltransferases were shown to be upregulated under leaf rust infection. At the adult stage, TraesCS1D02G004600, encoding a cytochrome P450, was expressed in the flag leaf blade at both the dough and ripening stages. Different studies have reported the role played by cytochrome P450 in the host response to disease, which included the response to Fusarium head blight (FHB) disease in wheat (Walter et al. 2008, 2011). The pathogen-responsive gene encoding cytochrome P450 has been shown to be differentially expressed under leaf rust infection in wheat (Bolton et al. 2008). Additionally, Bolton et al. (2008) reported that gene models coding for the same protein as some of the hcmQTLs discovered in this study were upregulated under leaf infection. All gene models coding for serine-threonine protein kinases and cytochrome P450 were upregulated in all treatments.

### Breeding implications for leaf rust resistance

The primary use of MQTLs for breeding purposes is the development of improved varieties with enhanced yield that are resistant to diseases via marker-assisted selection (MAS). MQTLs with the smallest confidence interval (CIs) have been harnessed effectively for MAS because they incorporate multiple QTLs, as reported for disease resistance in maize (Xiang et al. 2012; Wang et al. 2016), grain yield-associated traits in rice (Wu et al. 2016; Carrijo et al. 2017), seed quality in soybean (Qi et al. 2017) and anthesis time in wheat (Griffiths et al. 2009). To this end, the smQTLs were refined to 15 hcmQTLs, each of them incorporating at least 5 original QTLs and having a physical interval lower than 20 Mb and a genetic interval lower than 10 cM. In addition, meta-QTL analysis can be used to identify regions that confer resistance to more than one disease, and the marker information can be used for MAS (Ali et al. 2013). In this study, the hcmQTLs1B.4 region was identified to confer resistance to leaf and stripe rusts, thus making it a potential region to exploit for multiple disease resistance in wheat. Furthermore, breeding for durable resistance is desired in major breeding programmes. Durable resistance remains effective against a pathogen for a significant number of years (Johnson, 1981; Johnson, 1984). The combination of seedling resistance and adult plant resistance has been proven to confer prolonged resistance over several years (Kolmer and Oelke, 2006). In addition, various studies have ascribed durable leaf resistance to adult plant resistance rather than to seedling resistance (Figlan et al. 2020). Therefore, hcmQTL1D.2 discovered in this study can be harnessed to confer durable resistance in wheat, as it incorporates genes conferring both seedling and adult plant resistance. Another useful approach that could be harnessed in breeding for leaf rust resistance in wheat is gene pyramiding. Gene pyramiding involves incorporating multiple desired genes into a single variety. Gene pyramiding is broadly acknowledged by breeders, plant pathologists and farmers to improve disease resistance in wheat (Chen and Kang, 2017). A major requirement for gene pyramiding is to identify various QTLs or genes conferring resistance and then incorporate them into a high-yielding cultivar (Singh 1992). In several instances, this technique has been utilized in crops. For instance, long-term resistance was conferred when diverse genes were pyramided with leaf rust genes (Kolmer, 1996; Bhawar et al. 2011; Aboukhaddour et al. 2020; Babu et al. 2020). Additionally, in barley, MAS combined with gene pyramiding has been used to introgress resistance loci against stripe rust into numerous lines (Toojinda et al. 1998, 2000; Castro et al. 2003a, b; Richardson et al. 2006). To this end, the hcmQTLs discovered in this study can be utilized and exploited for gene pyramiding via MAS to bolster the resistance of wheat against leaf rust.

### Concluding remarks

One of the most effective methods for analysing the wealth of QTL information available from various studies is MQTL analysis. In this study, we delineated the genetic architecture of leaf rust in wheat via MQTL analysis and by integrating genomics studies. In comparison with initial QTL reports, meta-analysis allowed us to reduce the MQTL CI, thereby facilitating the search for candidate resistance genes in the databases available. The result was the discovery of 15 hcmQTLs, with each having a potential role in MAS. This result will be useful for developing resistance to leaf rust through the introgression of desirable hcmQTLs that could confer a high level of resistance during cultivar development. Last, this study can also help better define the various mechanisms associated with leaf rust resistance in wheat.

## Supporting information

Online Resource 1

Online Resource 2

Online Resource 3

Online Resource 4

Online Resource 5

Online Resource 6

Online Resource 7

Online Resource 8

Online Resource 9

Online Resource 10

## Acknowledgements

JMS acknowledges the contribution of the CERCA Program (Generalitat de Catalunya).

## Online Resources

ESM1: List of publications on past QTL mapping studies used for our meta-QTL analysis

ESM2: List of leaf rust resistance QTL identified in the mapping populations for meta-QTL analysis

ESM3: Leaf rust association studies used for MQTL validation

ESM4: Validation of smQTLs from previous GWAS on leaf rust in wheat

ESM5: Co-localization of leaf rust genes with smQTLs

ESM6: Discovery of candidate genes within the interval of hcmQTLs

ESM7: Differentially expressed genes from ERP013983

ESM8: Differentially expressed genes from SRP041017

ESM9: Differentially expressed genes from ERP009837

ESM10: Gene ontology for genes expressed within the hcmQTLs interval

## Notes

**Conflicts of interest/Competing interests** Authors declare no conflict of interest.

### Competing Interest Statement

The authors have declared no competing interest.

